# Investigating the Inhibitory Aspects of Metformin/Curcumin Co-Treatment through Convergence of In-Silico and In-Vitro Approaches

**DOI:** 10.1101/568634

**Authors:** Farzaneh Afzali, Zahra Nayeri, Zarrin Minuchehr, Mossa Gardaneh

## Abstract

Nearly 16% of people with breast cancer (BC) have Diabetes Mellitus type 2 (DM2) and are at a higher risk of death worldwide. Their common regulatory factors and functional mechanisms can be targeted applying multi-target drugs including Metformin (MTFN) and Curcumin (CURC). In this study, we used in-silico approaches to study the potential underlying mechanisms of this co-treatment strategy on BC and DM2 in order to introduce novel therapeutic targets.

The total number of 48 shared differentially expressed genes (17 up-regulated and 31 down-regulated) were identified through establishing diseases’ protein-protein network and BC RNA-sequencing expression data. The integration of functional clustering and pathway analyses revealed that the most involved cellular pathways and processes are regard to cells’ proliferation, death, migration, and response to external stimulus. Afterwards, the MTFN/CURC correlation and co-treatment optimization was probed through response surface methodology (RSM) based on MCF7 cell line and confirmed by MDA-MB-231. Combination index calculation by MTT viability assay proved supportive effects on both cell lines. The superior apoptotic potential of co-treatment compared to single treatments was shown on inhibition of MCF7 proliferation and induction of cell death demonstrated by cell body co-staining and flow cytometry as well as gene expression analysis via RT-PCR. Furthermore, wound-healing scratch assay showed that this co-treatment has a slightly higher effect on migration inhibition compared to single treatments.

In conclusion, our study used in-silico and in-vitro approaches and introduced a potential regulatory panel between BC and DM2. We also provided a linear model and equation that show the positive relation of drugs’ co-treatment. The proposed co-treatment strategy successfully controlled the biological processes under investigation.

## INTRODUCTION

Once disease causes are identified, solutions for disease management and therapy become readily traceable. Therefore, in the context of multi-factorial diseases, multi-target drugs based on targeting either multiple factors or conditions are amplified. Such multi-target strategies are indeed more beneficial to patients than single-target procedures for heterogeneous diseases like diabetes mellitus type 2 (DM2) or breast cancer (BC) that are also influenced by global factors like environment or lifestyle [1]. Multi-factorial mechanisms of disease development involve multi-genes complexes that are best functional within their appropriate biological networks including any related pathways, processes, functions, and even their niche of existence [2]. Metformin (N’,N’-dimethyl biguanide hydrochloride; MTFN) and curcumin (1,7-bis(4-hydroxy-3-methoxyphenyl)-1,6-heptadiene-3,5-dione; CURC) both have inhibitory effects on DM2 and BC, a criterion needed for investigating their underlying genes and mechanisms responsible for inhibiting these conditions.

BC and DM2 fuel one another’s occurrence considering their common underlying pathways of glucose metabolism. The impaired metabolism of abundant glucose and its related cascades constitutes the basis of the “Warburg effect” theory [3]. The Warburg effect states that, despite bioavailability of oxygen for effective ATP production via oxidative phosphorylation, cancer cells initiate less-productive glycolysis to produce their required energy fuel. Glycolysis does not produce enough ATP but the cell experiences up-regulation of genes responsible for absorbing more glucose known as metabolic reprogramming [4,5]. Metabolic reprogramming plays critical roles particularly in adaptation of cancer cells to restrictive conditions dictated by tumor microenvironment. The Warburg effect is triggered by up-regulation of molecules such as GLUT proteins that promote glucose transfer across cell membrane and accumulation of cytosolic glucose [6].

As an anti-DM2 drug, MTFN suppresses hyperinsulinemia and hyperglycemia by counteracting insulin resistance in DM2. MTFN inhibits gluconeogenesis and so glucose absorption from the small intestine and increases glucose uptake in the cells and reduces plasma free fatty acid concentration [7]. MTFN also increases insulin induced translocation of glucose transporters to the cell membrane and so reduces insulin resistance [6]. The anti-mitogenic effects of MTFN on cancer cells are exerted either by activating intracellular energy sensor adenosine monophosphate-activated protein kinase (AMPK) or via AMPK-independent signaling pathways [8]. Major impacts of MTFN on cancer include sensitization of resistant tumors to chemo-radio- and antibody-therapy [9–11], which are mainly caused by selective killing of tumor-initiating cells or cancer stem cells (CSCs) [12]. Such resensitizing potential of MTFN has been shown to improve the prognosis of diabetic patients who carry HER2+ BC tumors [13]. Combined with cancer drugs such as cisplatin or doxorubicin, MTFN indeed enhances their cytotoxicity and such combination synergistically reduces tumor size [14], an unexplored potential of MTFN that is being examined in clinical trials [15].

Curcumin (CURC) induces cancer cell death by a long list of inhibitory and inducing activities on cellular pathways involved in apoptosis, autophagy and growth inhibition [16]. Reduced cell growth, arrested cell cycle and diminished tumor size and inhibited metastasis are effects of CURC on HER2+ BC xenografts [17]. Our own study indicates that CURC can synergize with TZMB as well as with anti-oxidant enzyme glutathione peroxidase-1 in inducing apoptosis in HER2-overexpressing BC cells, indicating CURC potential to challenge TZMB [18]. Furthermore, Kakarala et al have demonstrated that CURC can be used to target CSCs in BC [19]. On the other hand, CURC affects glucose metabolism by inhibiting aerobic glycolysis in cancer cells [20].

The aim of the current study is to explore fortifying effects of MTFN/CURC co-treatment on inhibiting the redundant processes and genes achievable by *in-silico* studies and establish a statistical relation between them. The intermutual elements for BC and DM2 were identified by analyzing their protein-protein interaction (PPI) network and merging with RNA-sequencing expression data of BC. The enrichment analysis and functional annotation clustering were also carried out to recognize the underlying mechanisms of action. Our next step was to adjust MTFN/CURC co-effects on MCF7 cell line for establishing the statistical relation of drugs and then check whether this model is applicable to another BC cell line, MDA-MB-231. The type of induced cell death was also investigated via fluorescence co-staining, flow cytometry, and reverse transcription (RT) PCR. Phosphatase and tensin homolog (PTEN), which is down-regulated upon MTFN treatment, is a tumor suppressor gene that antagonizes the PI3K/Akt/mTOR [21,22]. If occurs, up-regulation of PTEN expression could be another anti-cancer dimension of MTFN/CURC co-treatment and diminishing their high-dose usage in single treatments.

## METHODS

### 1. Identification and validation of overlapped DEGs between BC and DM2

The STRING disease network importer of Cytoscape 3.6 was used to establish the PPI networks of BC and DM2 with ultimate number of proteins (2000 nodes) and 0.7 as the minimum interaction score. The networks were merged together to find the overlapped factors and subsequently, filtered by BC differentially expressed genes (DEGs) of The Cancer Genome Atlas (TCGA) database. TCGA database contains the RNA-sequencing (RNA-seq) expression data of 33 different types of human cancers. In this study, BC RNA-seq raw data of 224 samples (112 cancerous tissues and 112 adjacent normal ones) was downloaded and normalized using TCGAbiolinks package of R v3.5.2 software. To identify the (DEGs) the criteria of FDR < 0.05 and |logFC| > 1 were applied.

### 2. Functional Annotation Clustering and Pathway Analysis

To understand the biological meaning behind the obtained DEGs in groups, Functional Annotation Clustering tool of David 6.7 database, based on Kappa statistics and fuzzy heuristic clustering algorithms, was used to make the ontology report easier to follow [23]. Each cluster contains Gene Ontology (GO) terms including Molecular Function (MF), Biological Processes (BP), and Cellular Compartment (CC). The Group Enrichment Score is a −log mean of the members’ p-values to signify their general significance. The classification stringency was set to high, while other parameters were left unchanged. In parallel, GO and pathway analyses were also performed to have better control on classifying clusters for designing subsequent in-vitro experiments.

### 3. Cell Culture

Human BC cell lines MCF7 (Michigan Cancer Foundation-7) and MDA-MB-231 (Michigan Cancer Foundation-7) were cultured in Dulbecco’s modified eagle’s medium (DMEM) plus 10% fetal calf serum (FBS), 100 U/mL penicillin, and 100 μg/mL streptomycin and incubated at 37 □C, 5 % CO2. MTFN and CURC were both purchased from Sigma-Aldrich and dissolved in DMSO to form treatment stocks.

### 4. MTT viability assay

The MMT assay was carried out as we have reported [24]. Serial concentrations of MTFN ranging from 5 to 21 mM and of CURC in the range of 5-21 μM were applied; DMSO (drugs’ solvent) and untreated cells were used in parallel as positive and negative controls, respectively. The cells were incubated for 24 hours before incubating with the MTT solution in shaker for 30 min and reading by an ELISA reader at 580 nm wavelength. The calculated IC50 represents the treatment concentration that inhibits 50% of cells’ growth versus controls.

### 5. Co-Treatment Experiment Design and Measurement of Synergism and Antagonism

The Design Expert version 7.0.0 software was used to design and optimize the contribution of two factors (Xi = drugs) in combination applying response surface methodology (RSM) [25]. Regarding to the central composite rotatable design (CCRD), which enables us to predict and examine the effects of independent variables (drugs) on each other and dependent variable (response; survival in this study) at five levels, 13 different runs were calculated to determine the optimal doses of drugs in combination; it included 5 runs at center point. Considering narrowing down the drugs’ single treatment doses after replicating the MTT experiment, 9-13 mM for MTFN and 5-6 μM for CURC were applied as limits with Face Centered as the α-coefficient and ±0.5 additions --in line with keeping lack of fit (LOF) to insignificant-- made the floating numbers to integer at some cases after calculations’ designation.

We then calculated combination index (CI) for the two compounds using formula below and as we have described before [18]: CI = (D_1_/Dx_1_) + (D_2_/Dx_2_) + (D_1_D_2_/Dx_l_Dx_2_), where Dx_1_ and Dx_2_ represent the doses of MTFN and CURC alone needed to produce x percentage effect, and D_1_ and D_2_ indicate the doses of MTFN and CURC in combination required to produce this effect.

### 6. Evaluation of Cell Death by Acridine Orange/Ethidium Bromide (AO/EB) Co-staining and Flow Cytometry Annexin V-FITC/PI Double Labeling

Cells were co-stained according to our previous reports [26]. We applied the IC_50_ of MTFN, CURC and their combination as inhibitory treatments. The images of stained cells were captured using fluorescent microscope.

MCF7 cells were seeded 2 × 10^5^ per well in a 6 well plate. After treatments, the cells were trypsinized, were resuspended and incubated with the stains for 5 min at room temperature before being analyzed by a FACSCalibur at 488 nm wavelength. A total of 10,000 cells were used per sample and the data was analyzed in FlowJo 7.6.1 software.

### 7. RNA extraction, cDNA synthesis, and Reverse Transcription (RT) PCR

Total RNA was extracted from BC cell lines using RNX-plus Solution (CinnaGen, Tehran, Iran) according to manufacturer’s instructions. Purity and integrity of RNA were determined using Nano Drop ND-2000 spectrophotometer (Thermo Fisher Scientific, Wilmington, USA). Then, cDNA was synthesized using Maxime™ RT PreMix (Oligo (dT)15 Primer) (Daejon, South Korea) and RT-PCR analysis was carried out as previously described [24]. The primer pairs were used as follows: *GAPDH (Control)*: 5’-CCCCCAATGTATCCGTTGTG-3’ and 5’-TAGCCCAGGATGCCCTTTAGT-3’ (Accession: NG_007073); *BAX*: 5’-TGG AGCTGCAGAGGATGATTG-3’ and 5’-GAAGTTGCCGTCAGAAAACATG-3’ (Accession: NG_012191); *BCL2*: 5’-CTGCACCTGACGCCCTTCACC-3’ and 5’-CACATGACCCCACCGAACTCAAAGA-3’ (Accession: NG_009361), and *PTEN*: 5’-CGAACTGGTGTAATGATATGT-3’ and 5’-CATGAACTTGTCTTCCCGT-3’ (Accession: NG_007466). The reaction products were then visualized using Gel Electrophoresis and quantified using ImageJ1.51 software [27].

### 8. Wound Healing/ Scratch Assay

One day after seeding 3 × 10^5^ MCF7 cells on each well of a 6-well plate, a scratch was made by a micropipette yellow tip and the cells were washed and re-fed with serum-free medium. The Cells’ images were taken before and after treatment period and were analyzed using ImageJ software. The area between two sides of each wound was measured for monitoring cell migration.

### 9. Statistical Analysis

The experimental data were analyzed by GraphPad Prism 6 and was provided as mean ± standard deviation (SD) of two independent experiments. The significance of differences between groups was analyzed by Two-Way ANOVA. We considered *p*-value less than 0.05 as significant, and *p*-value less than 0.01 or 0.001 as highly significant.

## RESULTS

### 1. Validated overlapped DEGs between BC and DM2

The established PPI network of BC and DM2 embodied 2000 nodes with 26413 edges and 1665 nodes with 16035 edges, respectively; total number of 469 nodes with 5323 edges were attained by merging the corresponding networks. After filtering the obtained nodes with 1209 BC DEGs (510 up-regulated and 699 down-regulated) of TCGA RNA-seq data, validated amount of 48 nodes (17 over-expressed and 31 down-regulated) with 96 interactions were remained (Fig. 1).

**Figure 1.**
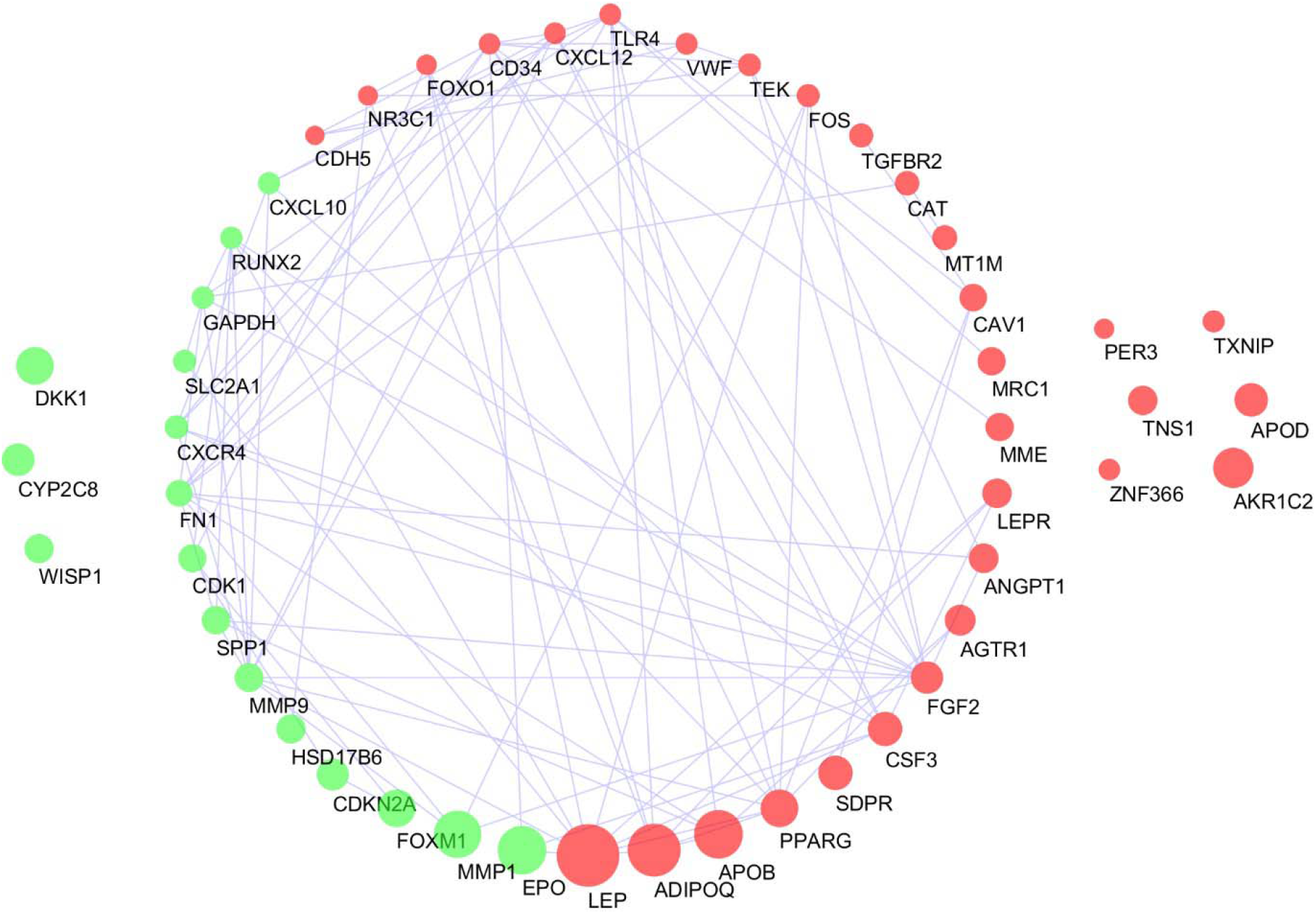
The PPI network of overlapped differentially expressed genes between BC and DM2. The network was provided and visualized by STRING database and Cytoscape software, respectively. The green and red nodes represent up- and down-regulated DEGs, jointly. The size of nodes indicate the intensity of fold change; bigger nodes are expressed more differently than the small ones.

### 2. Fundamental Annotated Biological Processes for In-Vitro Experiments

The remained 48 nodes were subjected to Functional Clustering Annotation tool of DAVID 6.7 database and 53 clusters were identified harboring corresponding enrichment scores (the higher is more enriched) and p-values. We added another column to the calculated clusters called average of p-values to find the significant groups (p-value < 0.05); this add-in filtered out 26 clusters. By virtue of comparing GO analysis and Functional Clustering results, it was found that cell migration, cell growth and death, response to external stimulus (hormones and drugs), and regulation of metabolic cascades are the most enriched cellular processes underlying overlapped genes between DM2 and BC. As it was expected, the genes were more concentrated in extracellular regions and were accountable for growth factors’ activities and elements’ bindings. The top five enriched pathways for common genes were Pathways in cancer, Cytokine-cytokine receptor interaction, Adipocytokine signaling pathway, Hematopoietic cell lineage, and Toll-like receptor signaling pathway. The summary of obtained data is available in Table 1 (for complete results refer to supplementary 1).

**Table 1.**
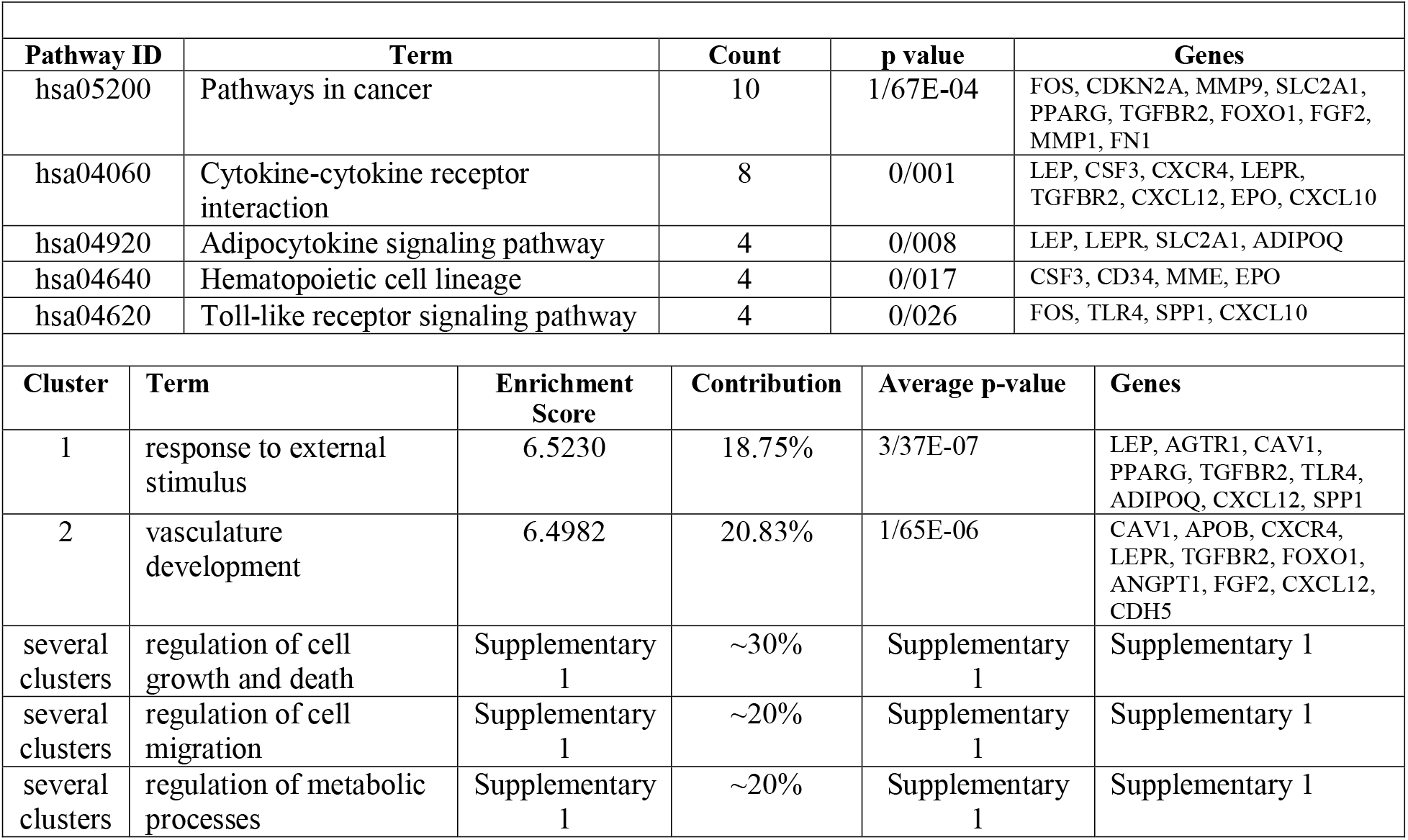
The enriched pathways and processes for overlapped genes between BC and DM2. The contribution column indicates the percentage of genes involvement in introduced terms.

### 3. Linear Model Validated synergic anti-proliferative effect of MTFN/CURC by MTT Viability Assay

According to CCD of RSM, five levels of combinations with limits are available in Supplementary 2. The obtained linear formula revealed one solution for reaching optimal (targeted) survival rate (50%) under MTFN/CURC co-treatment. The ANOVA analysis of the model provided a panel including F-value, p-value, Mean and sum squares, coefficient estimate that elucidates the synergic relation of variables in case of being positive and vice versa, and etc., which are available in Table 2. The p-value less than < 0.0001 and F-value of 36.85 suggested the linear model to be the appropriate predictor of response in the current study. In case of LOF, the F- and p-values were settled on 5.78 and 0.0892, respectively, indicating that this model can be fitted for our study. The R^2^ indicates the capability of model to calculate the amount of variation around the mean. Predicted and adjusted R^2^ should be within 0.20 of each other; the current model hold the difference of 0.0768 proving its reasonableness. In addition, the Adequate Precision value of 20.051 is greater than the cut-off of 4 that shows a discriminative model. The two- and three-dimensional plots of RSM and proposed equation of the model is available in Fig. 2.

**Figure 2.**
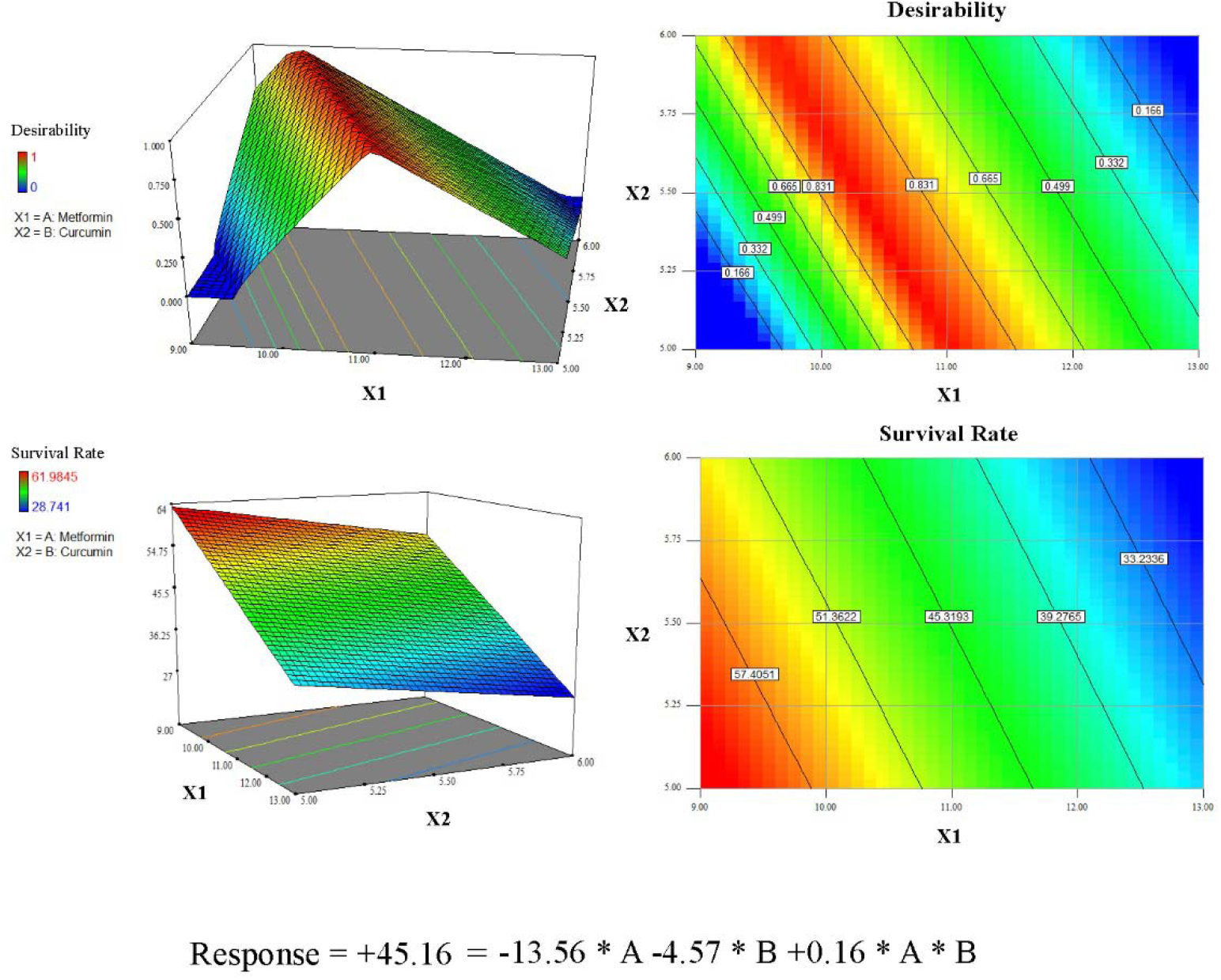
The plotted 2 and 3D graphs for introduced linear model of MRFN/CURC co-treatment. The graphs were plotted via RSM of Experiment Design software. The desirability plot shows the optimum combinations of drugs for reaching to 50 percent survival. The survival rate plot indicates the lower and upper limits of drugs’ combinations that was experimentally obtained by MTT assay. The provided linear equation is the mathematical formula of the co-treatment.

**Table 2.**
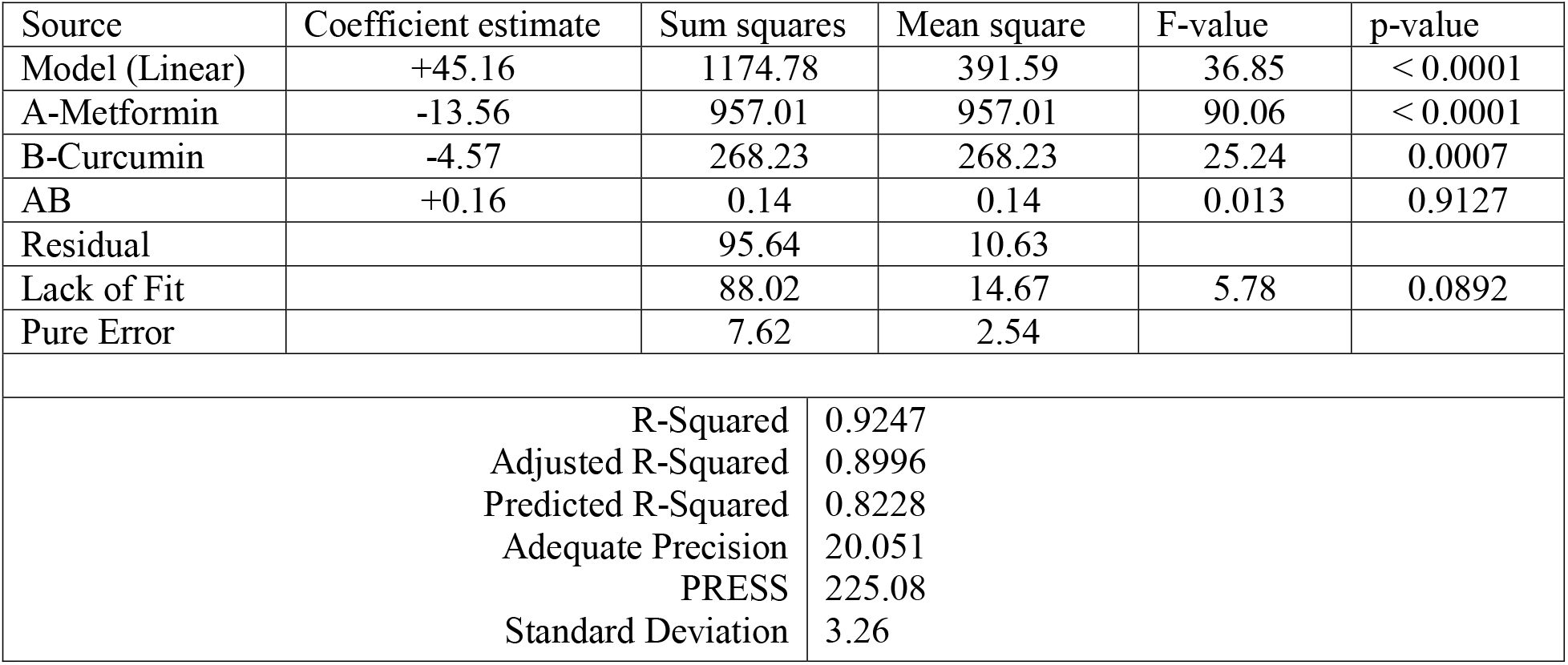
The ANOVA analysis of the suggested linear model and equation for survival rate of the MTFN/CURC co-treatment and its significance.

| Source | Coefficient estimate | Sum squares | Mean square | F-value | p-value |
| --- | --- | --- | --- | --- | --- |
| Model (Linear) | +45.16 | 1174.78 | 391.59 | 36.85 | < 0.0001 |
| A-Metformin | -13.56 | 957.01 | 957.01 | 90.06 | < 0.0001 |
| B-Curcumin | -4.57 | 268.23 | 268.23 | 25.24 | 0.0007 |
| AB | +0.16 | 0.14 | 0.14 | 0.013 | 0.9127 |
| Residual |  | 95.64 | 10.63 |  |  |
| Lack of Fit |  | 88.02 | 14.67 | 5.78 | 0.0892 |
| Pure Error |  | 7.62 | 2.54 |  |  |

|  |  |
| --- | --- |
| R-Squared | 0.9247 |
| Adjusted R-Squared | 0.8996 |
| Predicted R-Squared | 0.8228 |
| Adequate Precision | 20.051 |
| PRESS | 225.08 |
| Standard Deviation | 3.26 |

With reference to MTT assay, the IC_50_ of MTFN was 19 mM for MCF7 and 21 mM for MDA-MB-231; as for CURC, these values stood at 13 μM and 15 μM, respectively. The Two-Way ANOVA measured a significant difference (less than 0.001) between drug-treated cells and DMSO-treated controls. The Linear Regression analysis, which investigates and predicts the impact of one variable on another one, was performed on single treatments by Prism software. The R-square of MTFN and CURC on MCF7 and MDA-MB-231 were respectively, 0/9967, 0/9856, 0/9946, and 0/9934; the R^2^ is in a range of zero to one, where one is an optimum value for fitness of linear model. The slope deviation from zero were also significant (p-value < 0.0001). Deviation from linearity, which indicates that factors (drugs) interrelation is not linear, was measured subsequently; the p-values more than 0.05 proved that factors correlation is absolutely linear (Fig. 3a).

**Figure 3.**
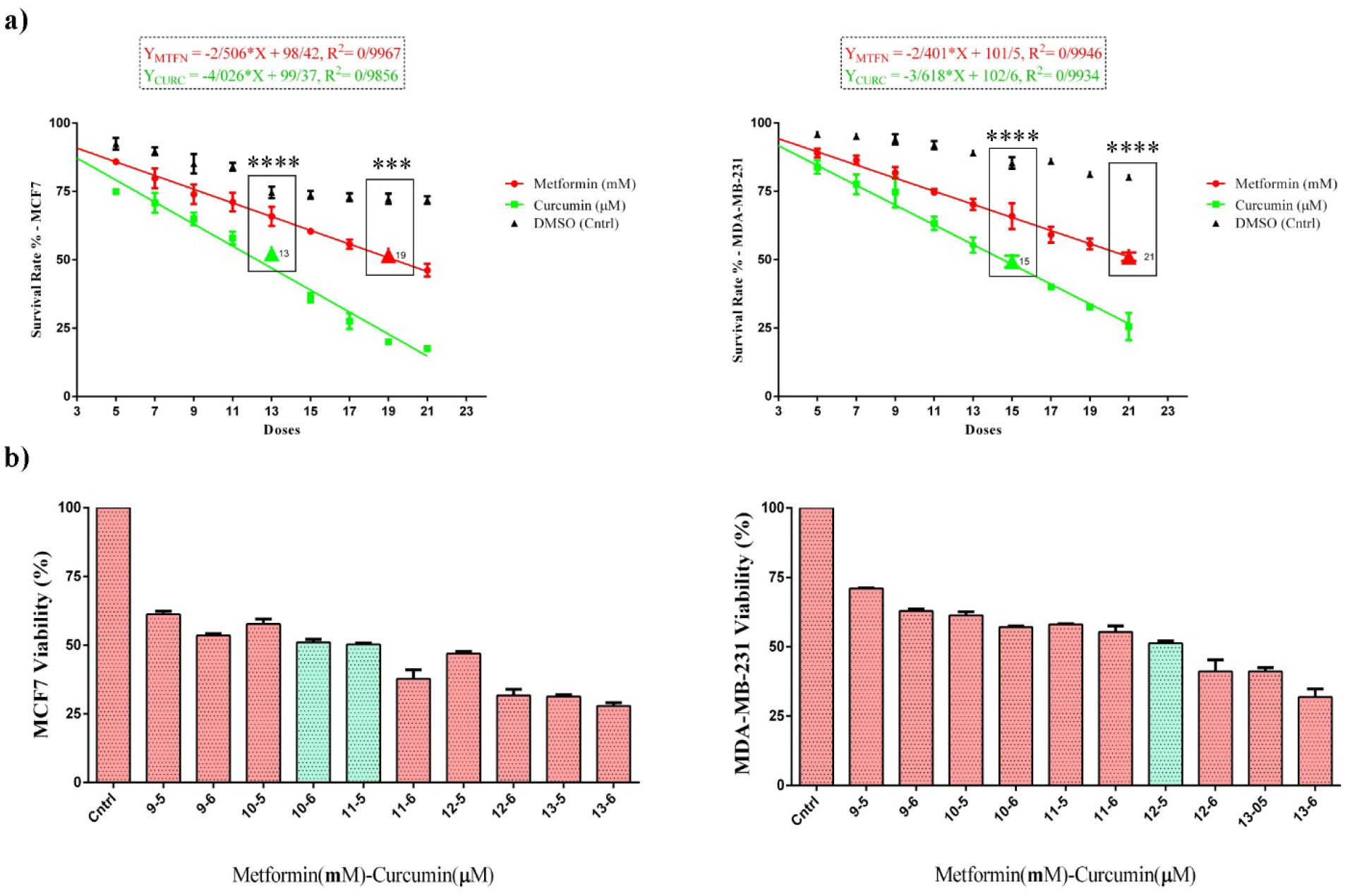
Survival plot of treated BC cell lines under MTFN, CURC, DMSO, and MTFN/CURC cotreatment by MTT assay. **a)** Each dot represents Statistical Reliability (SR) of two independent experiments with their error bars as standard deviation. The red, green, and black dots are MTFN (mM), CURC (μM), DMSO serial concentrations. The linear regression values and statistics that represent and confirm the linear interrelation of drugs are provided as lines, equation, and R-squared. The asterisk are result of Two-Way ANOVA showing the highly significance difference (p-value < 0.001 and 0.0001) of drugs and corresponding DMSO in terms of IC50 that embodied in black rectangles. **b)** The co-treatment survival plots (MTFN: 9-13mM and CURC: 5-6 μM). The cyan columns represent CI50 of co-treatment.

The CI50 value was calculated to better annotate the type of drugs’ correlation. It was found that cotreatment inhibits BC cells proliferation more than single treatments (Fig. 3b). The CI_50_ of MTFN and CURC in MCF7 and MDA-MB-231 were (10 mM - 6 μM or 11 mM −5 μM | CI ~ 0.96 = additive effect) and 12 mM - 5 μM (CI = 0.79= slight synergism).

### 4. Increased Growth Inhibition and Apoptotic Cell Death in Co-treated MCF7

To ensure the anti-proliferative or apoptotic nature of the observed reduced cell survival, AO/EB co-staining and flow cytometry were performed. Metformin-treated MCF7 cells exhibited proliferation inhibition and morphology changes from spindle shape to circular. Acridine orange/ethidium bromide (AO/EB) co-staining showed some yellow/green and partially orange cells as sign of apoptosis (Fig. 4a). CURC-treated dead cells outnumbered MTFN-treated ones. We also detected plenty of fully orange cells as sign of Necrosis. The number of cells remaining in co-treated wells was minimum indicating more profound effect of co-treatment on killing cells compared to single treatments.

**Figure 4.**
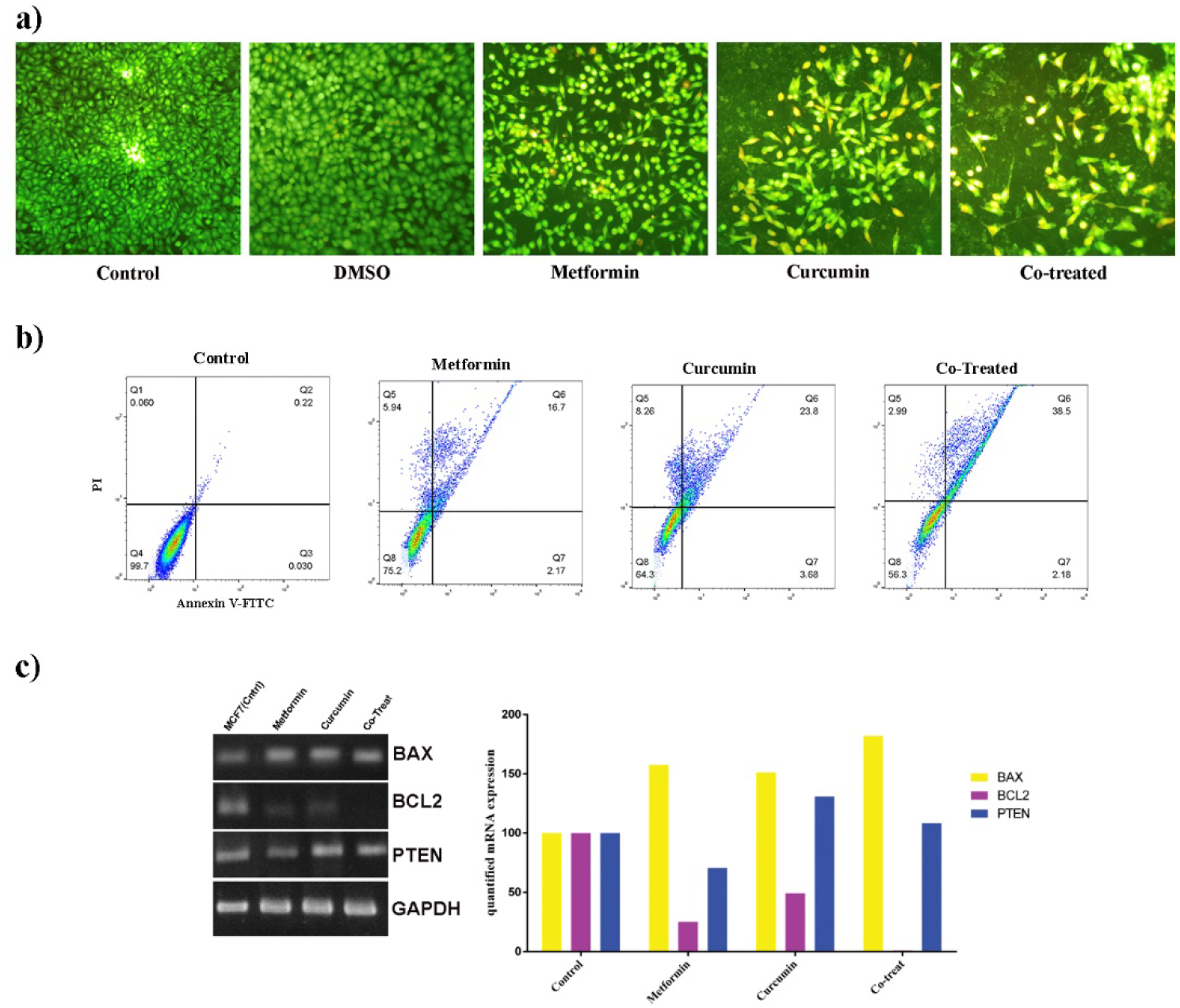
Examination of MTFN/CURC co-treatment induced growth inhibition and cell death in 24h exposure. **a)** Morphological changes of MCF7 under mentioned treatments by AO/EB co-staining. The cells were observed and were magnified 10X and 20X under fluorescent microscope. The green and yellow/orange cells represent alive and apoptotic ones, respectively. Apoptotic cells are more linear in shape rather than being circular. Lower density of cells under co-treatment towards control and single treatments indicates its strong potential in growth inhibition or cell death promotion. **b)** The MCF7 death evaluation by Annexin V-FITC/PI double labeling utilizing flow cytometry. The co-treatment is stronger in cell death induction in comparison to other experiments. **c)** Gene expression alterations examined by RT-PCR. The gel electrophoresis image represents expression of candidate mRNAs and the graphs were produced by quantification of each band within the image using the ImageJ software and plotted by GraphPad Prism 6. Each column represents SR of quantified mRNA expression.

Annexin V-FITC/PI double labeling fluorescence followed by flow cytometry was applied to more precisely monitor apoptosis among treated cells. As shown in Fig. 4b, CURC is more potent in killing cells so that it eradicated 35.74% of the treated BC cells compared to 24.81% killed by MTFN alone. Conversely, MTFN left 75.2% of cells alive whereas percent of live cells upon CURC treatment was 64.3%. The data demonstrate that proliferation inhibition is the dominant strategy MTFN adopts to stop cancer cell growth. We comprehended that co-treatment is more efficient against cancerous cells by inducing apoptosis (40.68%) more than single treatments.

Another way to ensure this finding is analyzing whether co-treatment alters apoptotic genes expression more than single treatments or not. As you see in Fig. 4c, both on electrophoresis gel and quantified chart, co-treatment significantly reduces expression of BCL2 as an anti-apoptotic gene and has slightly more effect on increasing expression of BAX as a pro-apoptotic gene. It also improves PTEN expression versus MTFN treated cells that is another proof for better effect of co-treatment in inducing anti-cancer effects because PTEN is a tumor suppressor.

### 5. Superior Effect of Co-treatment on Migration

Figure 5, demonstrates the pattern of BC cell migration in 24 h of being under the impact of applied drugs regarding the wound area. The result of these experiments showed that both single and co-treatments constrain cells’ movement, but co-treatment is a little more efficient rather than single treatments.

**Figure 5.**
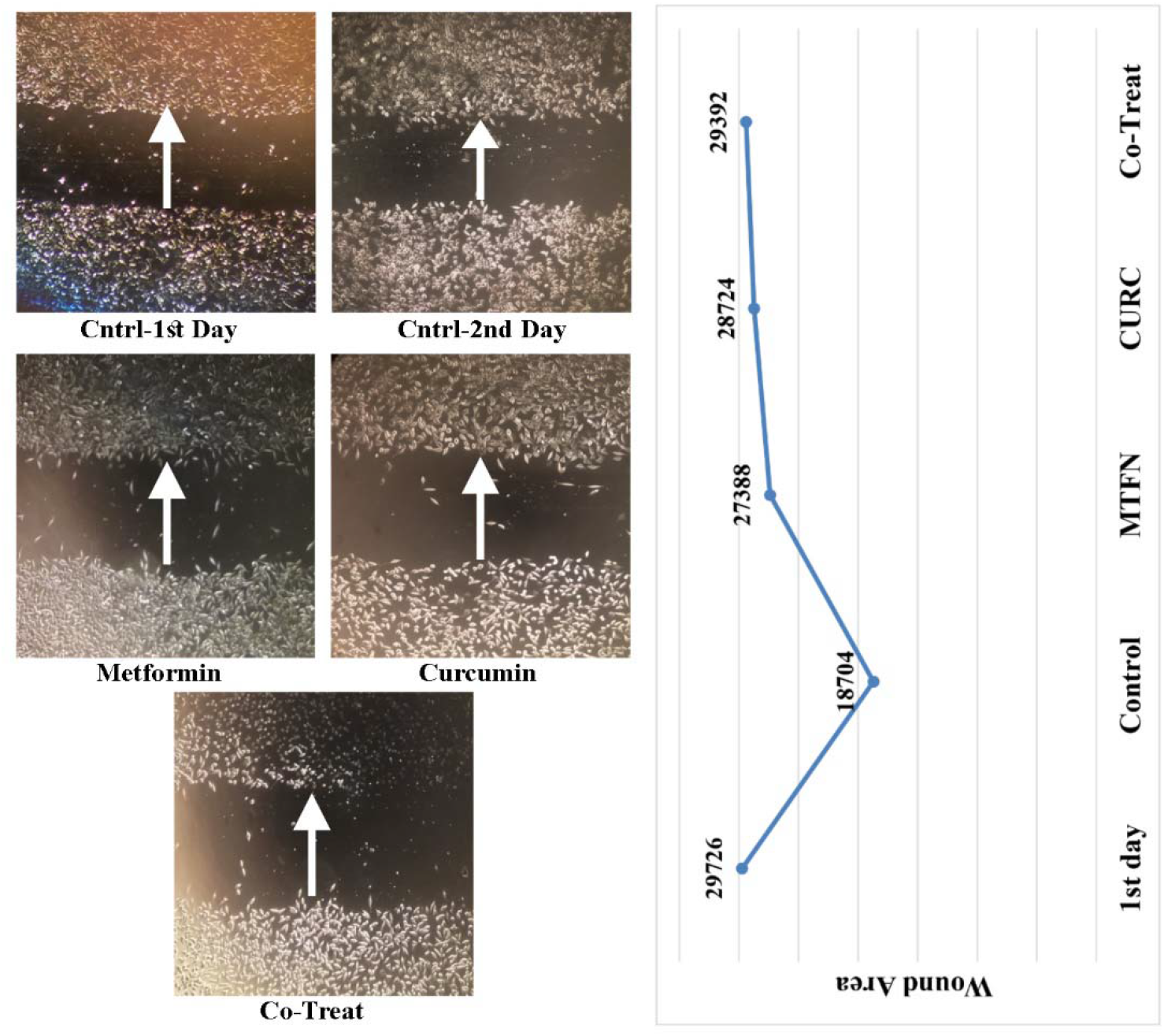
MCF7 wound-healing migration analysis. MCF7 migration was tested by wound-healing assay under IC50 of MTFN, CURC, and their combination after 24h. The cells were observed and magnified 4X under inverted phase contrast microscope. The quantified wound area was measured and expressed as the space between two sides of cultured cells by ImageJ software.

## DISCUSSION

In the current study, we mapped the common regulatory factors and mechanisms behind two human disorders BC and DM2 based on their sensitivity toward multi-target drugs, MTFN and CURC. These conditions possessed 469 common genes that were reduced to 48 ones after merging with expression RNA-seq data of BC TCGA samples. It was uncovered that these genes are generally responsible for regulating cell growth, death, and migration, hence are accountable for related pathways. The drugs’ correlation found to be linear utilizing RMSD methodology; this model was validated by MTT viability assay. The MTFN/CURC co-treatment exerted anti-proliferative effects, including both cell death and growth inhibition, on cancer cells that were manifested as slightly synergistic on MDA-MB-231 and additive on and MCF7 cell lines. The expression of BCL2 was decreased while that of BAX was increased under MTFN/CURC co-treatment compared to single treatments. Also, PTEN was overexpressed by co-treatment versus MTFN single treatment. Finally, the co-treatment promoted migration inhibition slightly more than single treatments.

Hard-to-cure human diseases such as diabetes, cancer or neurodegenerative disorders are often multifactorial in nature implying that a diverse range of underlying mechanisms behind disease genesis and progression need to be taken into consideration for improved management. An important manifestation of the biological networks is shown by gene regulatory networks that provide functional explanations for many biological phenomena and pathological conditions by identifying key molecules, their relevant pathways, and their causative relationships [28]. BC heterogeneity that drives tumorigenesis, metastasis, recurrence, and drug resistance necessitates drugs combinatory and multi-target strategies to be halted. The gene expression profiling, regulatory genes, and pathways involved in this phenomenon need to be taken to account for ultimate solutions [29]. Metabolic disorders like DM2 that result from desk-bound activities, also, are responsible for BC emergence and evolvement [30,31]. In DM2, insulin resistance or scarcity of insulin production enhances glucose level in blood [32] and as a sign of BC-DM2 adjoining, the extra glucose will be consumed by cancerous cells as their fuel to survive and grow. Together, MTFN and CURC appear to have converging points to synergize in halting tumorigenicity as a whole but also each can affect specific molecules or pathways within cancer cells. Co-treatment with MTFN and CURC indeed inhibits cancer growth, metastasis, angiogenesis *in-vitro* and *in vivo* [33], modulates immune system, and induces apoptosis independent from p53 in BC [34].

In virtue of using in-silico technics in the current study, one of the identified crosstalks that draws a link between BC and DM2 processes and pathways is the metabolic regulation accompanied by inflammatory responses and cytokines or hormones. Accumulation of metabolic hormones such as Leptin, producible by adipocytes, can cause hypoxia and in return can trigger growth, angiogenesis, inflammation, and immune reactions [3,35]; these factors can be controlled by MTFN and CURC single treatments [36,37], so by carrying out the MTFN/CURC co-treatment the chance of limiting the mentioned cellular processes would be increased. The correlated discovered genes through this study can be targeted by this co-treatment and keeping away the encouraging problems for both conditions.

Mitochondrial dysfunction involved in intrinsic apoptotic pathways relevant to tumor formation contributes to development of both BC and DM2 [38–40]. We and others have reported reduced BCL2 expression and increased BAX expression by CURC and MTFN single treatments [41,42]. Now, our current study reports that CURC-MTFN co-treatment more significantly induces BCL2 downregulation and BAX upregulation, hence inducing apoptosis more efficiently compared to single treatment. We found that MTFN adopts a strategy of inhibiting proliferation rather than inducing cell death. This may be due to MTFN ability to reduce circulating insulin leading to downregulation of proliferation pathways that are physiologically triggered by insulin or insulin growth factors [43]. The findings were quantified by AnnexinV-FITC/PI double labeling in flow cytometry that confirmed MTFN proliferation inhibition and co-treatment cell death induction.

PTEN plays complex roles in multiple cellular processes including proliferation, survival, apoptosis, cell cycle regulation, adhesion, and migration [21,44]. In our study, PTEN expression was reduced upon MTFN treatment, in line with previous studies [45,46], but this reduction was compensated following MTFN/CURC co-treatment. Cell death induction by co-treatment suggests that the co-presence of MTFN and CURC may strengthen the anti-tumor activities of PTEN via engaging several pathways and factors including miR-21 and Akt/PI3K [47]. This restriction may down-regulate or abate the insulin signaling pathway and result in insulin accumulation, insulin resistance, and hyperglycemia thereby co-promoting DM2 and BC. Therefore, PTEN reduction can help in the optimal use of insulin in cells [48,49]. MTFN promotes PTEN down-regulation through AMPK dependent mode [43] and beside its anti-DM2 effects, controls BC indirectly via the Warburg effect. The AMPK pathway acts as a sensor of energy in cells and further plays a major role in insulin signaling [50]. Cell metastasis is a major complexity that makes cancer hard to control. Both MTFN and CURC have anti-metastatic effect on cancer cells [51,52]. The wound healing assay showed more efficient but insignificant differences in cell migration in our MTFN/CURC co-treated samples compared to single treatments. MTFN and CURC effects on angiogenesis genes that help the migration phenomenon are controversial and cell-dependent especially in BC [53–57]. Therefore, further investigations are needed to monitor changes in the expression of specific genes in migration-related pathways so to understand the mechanisms associated with this co-treatment.

Previous studies like Zhang et al. [33], have examined the impact of MTFN/CURC co-treatment on BC, but the current study have inspected the underlying mechanisms and factors rooting from DM2 promoting PPI network and in parallel, designed a predictive statistical model for drugs’ correlation. It should be stated that this study may have multiple limitations that can be completed through future studies; the xenografts’ tumors derived from BC patients who suffer from DM2 as well, provide suitable in-vivo models for simultaneous evaluation of this co-treatment on both conditions. Also, the expression evaluation of involved factors in the introduced PPI network can be another phase of study to fully comprehend the underlying mechanisms.

### Conclusion

In conclusion, we have designed a linear model for MTFN/CURC interconnection based on MCF7 cell line and used MTT viability assay and confirmed its compatibility to another BC cell line, MDA-MB-231. Furthermore, the differentially expressed genes based on BC expression data were introduced that may have promoting potentials on DM2 as they were mapped upon merged networks of both diseases. The obtained processes were consistent with our in-vitro experiments showing that this inhibitory strategy is more effective than single treatments.

## Acknowledgment

This study was financially supported by a grant (502) from national institute of genetic engineering and biotechnology (NIGEB).

