## Supplementary 2 for "Investigating the Inhibitory Aspects of Metformin/Curcumin Co-Treatment through Convergence of In-Silico and In-Vitro Approaches"

**Supplementary Table 1.** RSM provided five levels of independent variables.

| Factor | Name | Unit | Level | | | | |
| --- | --- | --- | --- | --- | --- | --- | --- |
|  |  |  | -1 | +1 | Mean | -α | +α |
| A | Metformin | mM | 9 | 13 | 11.077 | 9 | 13 |
| B | Curcumin | μM | 5 | 6 | 5.462 | 5 | 6 |

**Supplementary Table 2.** The 13 Different runs and obtained result by MTT viability assay on MCF7 cell line.

| Run | Factor 1: A | Factor 2: B | Response 2 |
| --- | --- | --- | --- |
| 1 | 9.00 | 5.00 | 61.9845 |
| 2 | 13.00 | 5.00 | 30.8441 |
| 3 | 11.00 | 5.00 | 50.5995 |
| 4 | 11.00 | 5.00 | 48.5489 |
| 5 | 10.00 | 5.00 | 56.3549 |
| 6 | 10.00 | 6.00 | 51.7578 |
| 7 | 12.00 | 5.00 | 46.3333 |
| 8 | 12.00 | 6.00 | 33.2614 |
| 9 | 13.00 | 6.00 | 28.741 |
| 10 | 12.00 | 5.00 | 46.57 |
| 11 | 11.00 | 6.00 | 36.7359 |
| 12 | 9.00 | 6.00 | 52.9875 |
| 13 | 11.00 | 6.00 | 40.048 |

**Supplementary Table 3.** The point prediction and actual responses by MTT viability assay on MCF7 cell line.

| Factors | Solution | PI low | PI High | Predicted Response | Actual Response |
| --- | --- | --- | --- | --- | --- |
| A: 11 | A: 10.55 | 37.99 | 52.31 | 45.1527 | 47.90 to 50.59 |
| B: 5.5 | B: 5.5 |  |  |  |  |
